# Personalizing the control law of an upper-limb exoskeleton using EMG signal

**DOI:** 10.1101/2021.09.23.461504

**Authors:** Benjamin Treussart, Remi Caron, Franck Geffard, Frederic Marin, Nicolas Vignais

## Abstract

Implementing an intuitive control law for an upper-limb exoskeleton dedicated to force augmentation is a challenging issue in the field of human-robot collaboration. The goal of this study is to adapt an EMG-based control system to a user based on individual caracteristics. To this aim, a method has been designed to tune the parameters of control using objective criteria, improving user’s feedback. The user’s response time is used as an objective value to adapt the gain of the controller. The proposed approach was tested on 10 participants during a lifting task. Two different conditions have been used to control the exoskeleton: with a generic gain and with a personalized gain. EMG signals was captured on five muscles to evaluate the efficiency of the conditions and the user’s adaptation. Results showed a statistically significant reduction of mean muscle activity of the deltoid between the beginning and the end of each situation (28.6 ± 13.5% to 17.2 ± 7.3% of Relative Maximal Contraction for the generic gain and from 24.9 ± 8.5% to 18.0 ± 6.8% of Relative Maximal Contraction for the personalized gain). When focusing on the first assisted movements, the personalized gain induced a mean activity of the deltoïd significantly lower (29.0 ± 8.0% of Relative Maximal Contraction and 37.4 ± 9.5% of Relative Maximal Contraction, respectively). Subjective evaluation showed that the system with a personalized gain was perceived as more intuitive, and required less concentration when compared to the system with a generic gain.

## I. Introduction

### A. Context

Musculoskeletal disorders (MSDs) are conditions that can affect muscles, bones, and joints. The appearance of MSDs is favoured by straining or repetitive tasks, making industrial workers particularly exposed [1]. MSDs have become a major health issue, impacting worker’s integrity as well as economics, by being responsible of a loss of porductivity and high healthcare costs. It was estimated that, in 2012, in France, the average cost per case of MSD was 21 k € [2]. In 2017 MSD represented 87% of the occupational diseases [3]. MSDs could be prevented by relieving physically the workers during straining tasks like load carrying.

In this context, exoskeletons could become a promising solution for industrial load carrying. Various designs of exoskeleton have been proposed, for a wide range of applications. Good transparency is defined as a minimum loss of ernegy in friction when transmitting an effort from the end effector of a robot to the actuator. Transparency is a key points for exoskeleton applications. It can be achieved mechanically [4], or with a control system [5]. However, in the case of direct manipulation (where the human is the one handling the load), the exoskeleton needs to apply relevant forces on the segment of the human in order to relieve his physical strain. In that case, transparency is not enough, since it only ensures minimal interaction forces between the robot and the human. The critical point is thus being able to determine what are these “relevant forces”.

In order to augment body capabilities, user’s intentions have to be estimated. This process can be performed by using force/torque measurements [6], but it can prevent direct manipulation, as the measurement unit usually needs to be between the load and the user’s hand. Muscle activity measurements are a common solution that enables direct manipulation. It is usually done with electromyography (EMG) sensors placed over the skin. This method does not affect the handling of the load. Different approaches to process and exploit the EMG signals have been developped in order to control a robotic system. It can be based on biomechanical models [7], proportional mapping [8], or machine learning algorithms [9].

A critical point in augmenting body capabilities is to ensure stable and safe interactions between human and robot. It is all the more critical in the case of exoskeletons because of the tight kinematic link between robotic and human bodies. In the current study, the objective is to tune and test personalized parameters that enable an intuitive interaction for a specific control system [10].

### B. Previous Works

EMG signals have already been used to control robotic devices [11] ,[12]. EMG signals processing can be performed into two different manners: discretely or continuously. Discrete methods consist in pattern recognition based on handcrafted features and a classifier trained with them [13] or some end-to-end neural networks [14]. There is a trade-off between the panel of available actions on one hand and the extense of the training and precision of the classification, which can depend on the number of classes, on the other hand.

The second type of approaches of EMG processing are continuous methods. These methods are more flexible but they also present disadvantages. The relations between the features of the EMG signals and the movement are highly non-linear [15]. In [16], a continuous 3D estimation of the position of the hand is performed with the use of 9 electrodes targeting specific muscles. Such precise requirements in the placement of the electrodes are laborious and examiner-dependent. These methods are particularly suited to teleoperation or prosthesis assistance. However, their use to control an exoskeleton appears difficult given that the assistance provided by the exoskeleton and the EMG activity of the user influences one another.

Some works have focused on the combination of the two types of approaches in order to benefit from the advantages of both sides. In [17], a system was designed where the discrete component was used to recruit the most pertinent continuous subpart. This allowed to design continuous subparts more limited but simpler to calibrate which brings flexibility to the system. And a higher-level discrete component that can have a limited number of available classes, making it more reliable.

In a previous study, a system that permits to carry an unknown load has been presented [10]. The assistance was based on EMG signals, measured with an armband placed around the arm (biceps and triceps recordings). The system was based on an integral corrector, aiming to reduce muscle activity of arm muscles obtained from EMG specific processing [18].

The system presented and tested in [10] consists of an intention detection block feeding an integral corrector as shown on Fig. 1. The intention detection module is composed of two processes : (i) the estimation of the direction intended and (ii) and the estimation of the intensity of the movement intended. The intensity’s estimation uses a model based approach [19], [20]. The direction’s estimation exploits a convolutionnal neural network inspired from computer vision architectures [21], [14](cf. Fig. 2). A key difference with computer vision architectures is the use of 1D convolutions instead of 2D convolutions and they are shared over the eight sensors of the EMG armband in order to extract the same type of features. The method in [18] can be compared to [17] as it is a mixture of continuous and discrete components used to process the EMG signal.

**Fig. 1:**
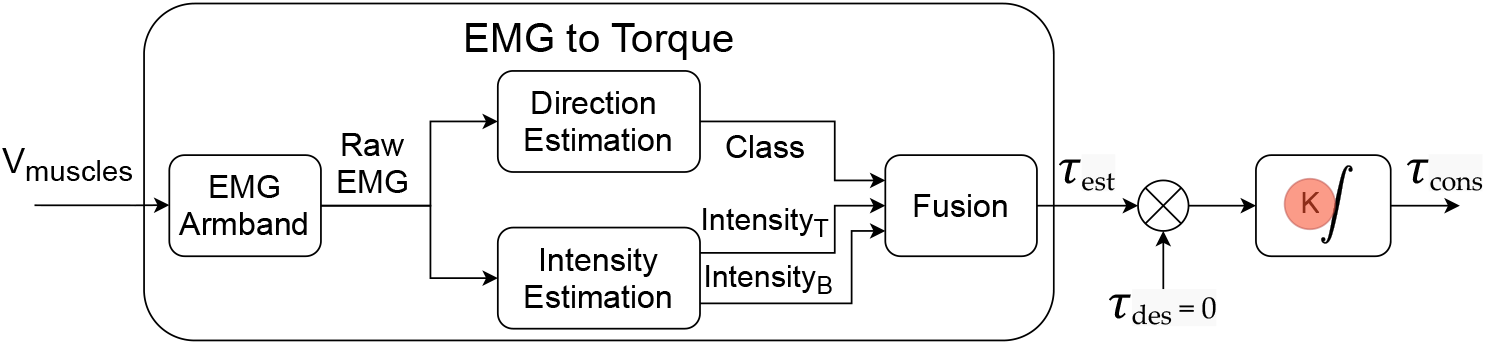
Intention Detection block and integral corrector from [10], K - the gain of the corrector.

**Fig. 2:**
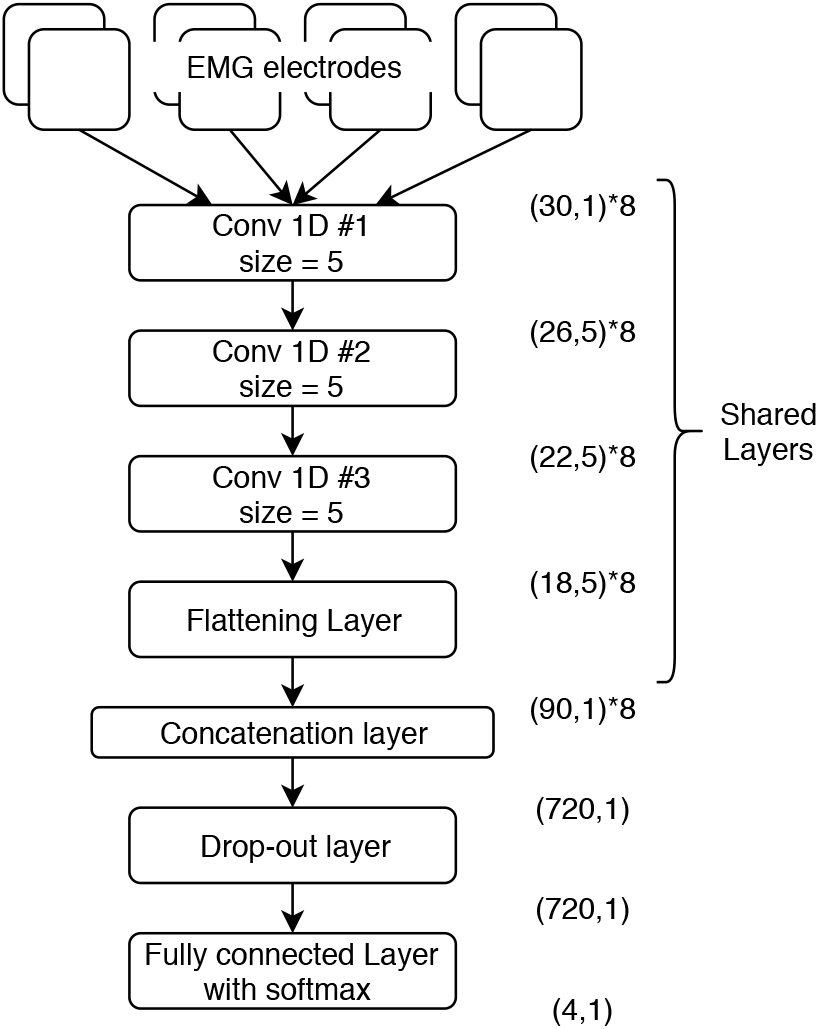
Architecture of the neural network

The system needed to be calibrated, as the EMG signals vary greatly from one person to another. The calibration data is required for both the calibration of the model based intensity estimation and the training of the network. In this study [18] the system is calibrated from scratch for each user, using about 2min30sec of recordings.

In the original study [10], the gain of the integrator is fixed. The system is compared to a classic gravity compensation (CG) and a situation without assistance (No-Exo). With a limited number of repetitions, results show (i) an EMG activity similar between the two types of assisted situations (CG and integral corrector) and (ii) a significant reduction of activity for the biceps, anterior deltoid and erector spinae when compared to the situation without assistance (integral corrector and No-Exo).

Our objective is to explore the possibilities of personalization regarding the tuning of the integrator’s gain, during a long duration, thus permitting to investigate user’s adaptation to the assistance conditions.

## II. Methodology

In the current study, the gain of the integratal corrector has been personalized using the user’s response time. Response time is defined here as the lapse of time between a stimulation and the response on the EMG signal. It is different from what is usually referred to as reaction time and reflex time [22] Reaction time is the delay between a stimulus and a reaction (ex: time to push the brake after spotting a pedestrian). A reflex time is the time for movement to occur after a stimulus on the muscle fibers (myotatic reflex). In our case the stimulation would be the increase of assistance from the system. Our hypothesis is that the user’s response time can influence the stability of the human/exoskeleton interaction (HEI). Personalizing the gain of the integrator would thus ensure a safe and intuitive interaction.

Our work is divided as follow :

1. Simulation to investigate the link between response time and gain of the integrator
2. Measuring response time on participants
3. Testing two control conditions through an experimental protocol: generic and personalized

The link between these different parts is described in Fig. 3. The first step of this process is to implement a simulation of the HEI loop. It enables to define the limiting values of the gain based on theoretical user’s response times. The results of the simulation give the relation between the gain of the corrector *K* and the user’s response time *Tps*_*rep*_. The second step consists in measuring actual user’s response times. Knowing that, and with the result of the first step a personalized gain *K*_*conf,new*_ can be estimated. The last step is the experimental protocol, its objective is to evaluate the impact of the personalized gain *K*_*conf,new*_ compared to a generic one *K*_*GEN*_.

**Fig. 3:**
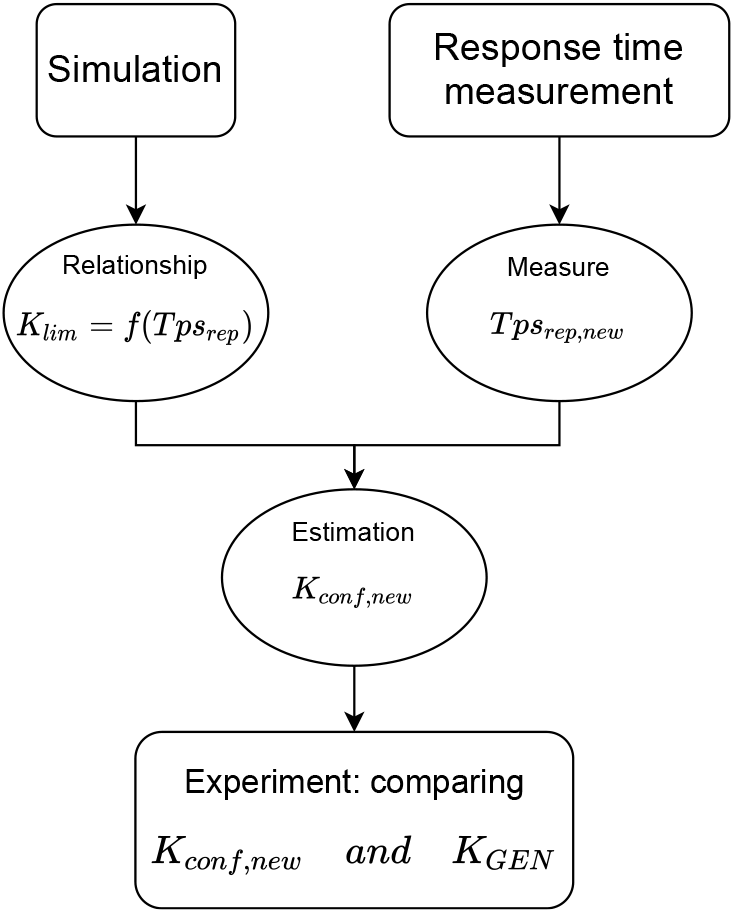
Flowchart of the study (variables are explained in table III)

As mentionned in section I-B, the system needs to be calibrated. In this study a shorter calibration procedure is used than in [18]. This version is based on a dataset created with the data of around 20 people recorded by following the procedure presented in [18]. This dataset is used to pretrain the neural network. A short step (15 secondes) of voluntary contractions is recorded in order to calibrate the system to a new user. The data of this step is used to calibrate the intensity’s model and fine-tune the network.

### A. Simulation

#### 1) Implementation

The interaction loop implemented is represented in Fig. 4 and the software used is Simulink®. The simulation is about a lifting task. In the beginning of a simulation, the load is at rest on a table and the user raises it up to a target. The system simulated has one degree of freedom, and is assimilated to a pendulum composed of a rigid arm and a load at the extremity. The real-life corresponding situation is a lifting task during which the elbow is kept straight. The objective of the simulation is to assess the relationship between a user’s response time and the gain of the integral corrector. This would enable to personalize the tuning of the gain and hopefully offer a more adapted behavior from the system of assistance.

**Fig. 4:**
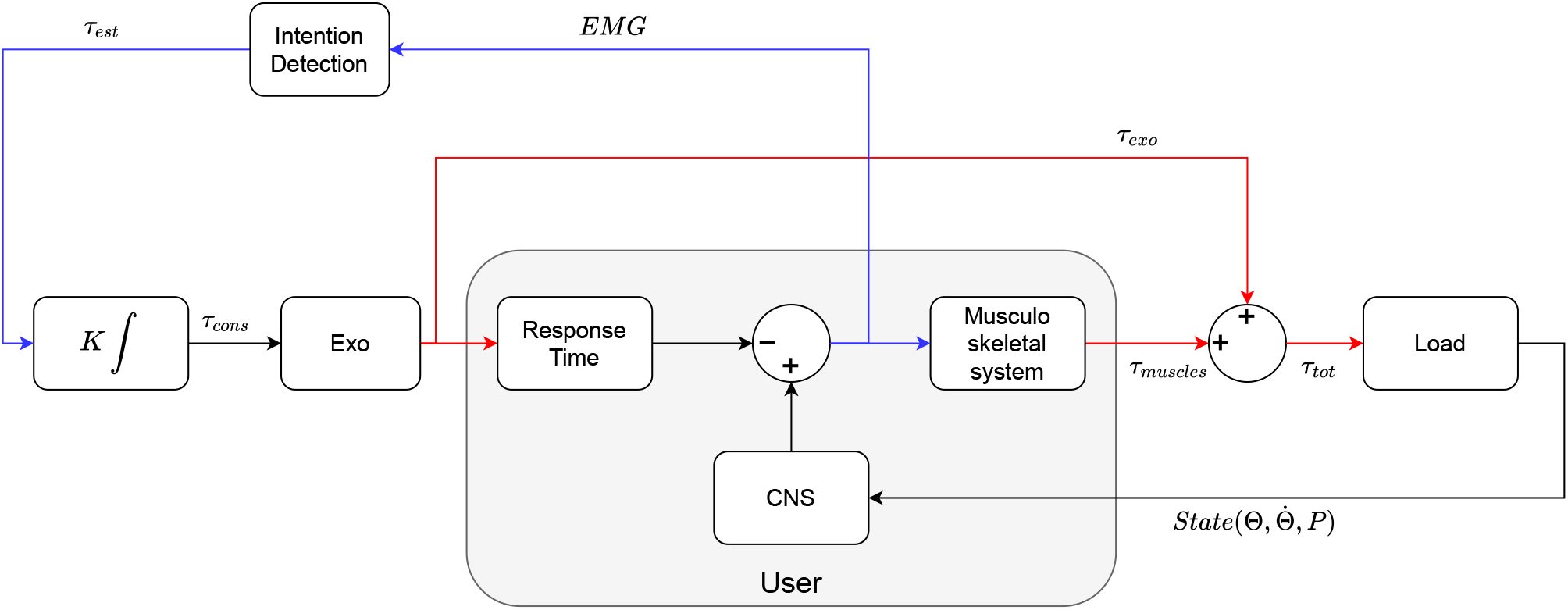
Interaction loop

##### a) User behavior Model

The implementation of the simulation of the user is divided into three parts. The first one is the response time. It is implemented as a pure delay, representing the time it takes for the user to perceive the changes of force from the robot. The second one represents the Central Nervous System (CNS), responsible of generating the setpoint signal to perform the lifting task.

The trajectory is defined using the minimum jerk theory [23], and studies have shown that it still holds while wearing an exoskeleton [24]. This theory enable to calculate a position Θ_*ref*_ (*t*) and velocity 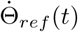 profile depending on the time. The total time for the movement was chosen according to the results of [10]. The output (cf. eq. (1) is given by the sum of a proportional-derivative (PD) controller, used to follow the trajectory, and the compensation of the weight of the load *τ*_*weight*_.

The perceived assistance provided by the robot is substracted from the setpoint given by the CNS. That is how the setpoint for muscle activation is calculated. It was considered that the user naturally adapts its effort according to the perceived assistance.

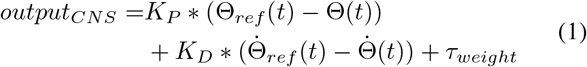

The force deployed by the user is modeled with the work presented by [25], that extends the Hill model with a Serial Damping Element in addition to the usual Contractive Element, Parallel Elastic Element and Serial Elastic Element.

The parameters are tuned using the data from the experiment of lifting task without assistance in [10], where there is an average activation of the anterior deltoïd 40% of Relative Maximal Contraction (RMC) for a load of 5kg.

##### b) Kinematics

During the interaction, human and robotic forearm dynamics are coupled, which can be modeled as follows in eq. (2) [24] :

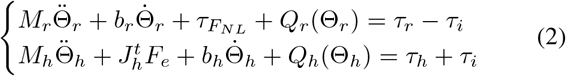

Where *τ*_*i*_ is the equivalent interaction torque between human and robot limbs, *τ*_*r*_ - the robot torque, *τ*_*h*_ - the human joint torque and *M* - the matrix of inertia, *b* - the viscosity, 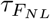 - the non-linear friction and *Q* the gravity torque. *F*_*e*_ are the external forces applied to the human and 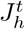 - the jacobian of the human. The subscripts *r* and *h* denote similar quantities related to robot and human systems, respectively.

Because only one degree of freedom is considered, we have *Q*_*r*_(Θ_*r*_) = *m*_*r*_*gl*_*r*_*cos*(Θ_*r*_), where *m*_*r*_ is the mass of the robot arm, *l*_*r*_ - the length to its center of mass and *g* the gravitational constant. The same is true for the human. The arm of the robot and the human are considered superimposed so Θ_*r*_ = Θ_*h*_. By combining that with the equation eq. (2), we have eq. (3):

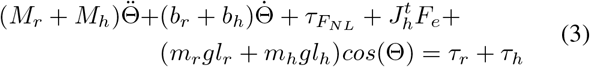

*τ*_*r*_ is divided as shown in eq. (4) :

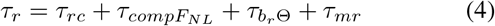

with 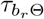, *τ*_*mr*_, and 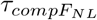 that can compensate 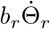, *m*_*r*_*gl*_*r*_*cos*(Θ_*r*_) and 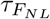, respectively, with classic control strategies, eventhough they can be very low due to the back-drivable mechanical design [26][4]. Taking this into account, and that the external forces come from the load *M* to be carried, the final equation eq. (5) is :

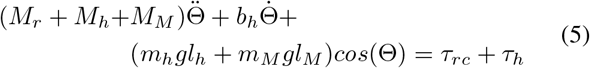

#### 2) Results

The simulation was exploited by making a search grid with the response time and the gain as parameters. The criterium to evaluate the simulation for a given pair of Response time and Gain (*Tps*_*rep*_, *K*) is the index presented in eq. (6). *K* varies from 4.5 to 8.2 with a step of 0.05 and *Tps*_*rep*_ - from 110ms to 210ms with a step of 5ms, these ranges are summarized in table **??**.

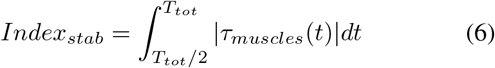

*T*_*tot*_*/*2 is set so that the system has enough time to stabilize if it is able. By design, if the system is stable the value of *τ*_*muscles*_ is close to zero. *Index*_*stab*_ is then an indicator of whether the system is able to stabilize. This index has been plotted on the Fig. 5. The yellow part corresponds to unstable simulations and the cliff is the limit between stable and unstable values of the parameters. The intersection of this surface with the plane *z* = 1000*N*.*m*.*s* gives the limit gain before instability *K*_*lim*_ as a function of *Tps*_*rep*_.

**Fig. 5:**
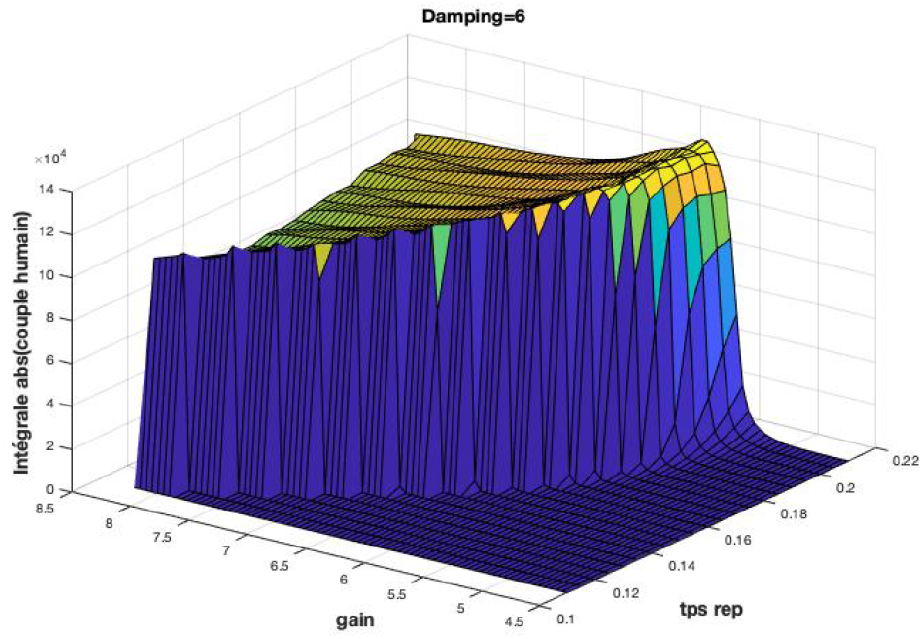
Value of the *Index*_*stab*_ for each pair (*Tps*_*rep*_, *K*) of the search grid

On Fig. 6 the intersection is plotted with a 2^*nd*^ order polynomial extrapolation. This extrapolation is used to estimate the *K*_*lim*_ of a new user based on their measured *Tps*_*rep*_. Except the *K*_*lim*_ is not the most practical to use, it tends to overshoot and cause slight oscillations before the system stabilizes. Instead, an expert tunes the gain *K*_*conf,ref*_, a comfort gain which oes not overshoot, and gets its *Tps*_*rep,ref*_ measured. The gain for a new user is calculated with eq. (7) :

**Fig. 6:**
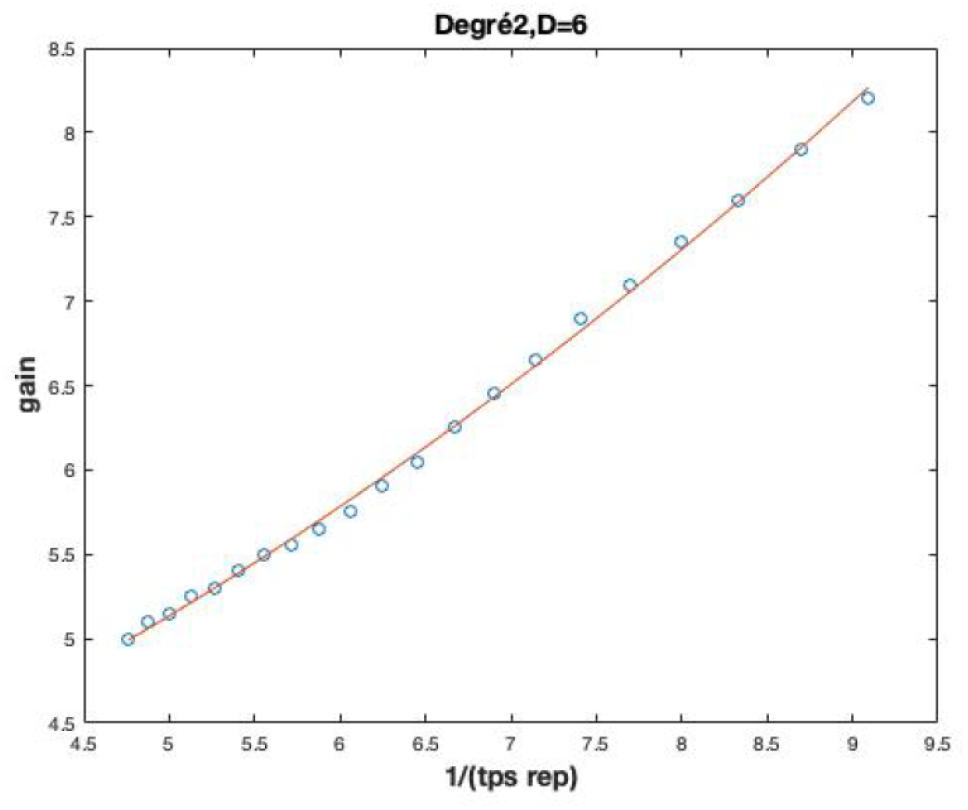
Graph of *K* _*lim*_ as a function of 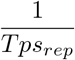, and a polynomial extrapolation of degree 2

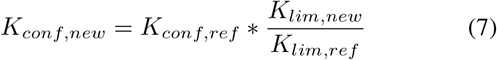

The simulation enables to estimate a range of possible gains that ensure the stability of the system, *K* ∈ [0, *K*_*lim*_]. The eq. (7) gives the possibility to take advantage of the experience of an advanced user and transfer it to a new user while taking into account their differences.

### B. Preliminary tests

Preliminary tests are conducted with a limited number of participants (5) in order to develop an automated way to measure the user’s response time. Participants are aksed to stay relaxed while jolts of torque are made with the exoskeleton (see Fig. 7). The jolts are triggered randomnly to avoid anticipation. For the preliminary tests, the jolts are applied in two different positions, high and low, and 10 times in each direction, for a total of 40 torque jolts. The objective is to find an indicator of the response time that does not vary greatly between the different conditions (direction and position) but still enables to discriminate between participants. The variables involved in this section are described at the end, in table III.

**Fig. 7:**
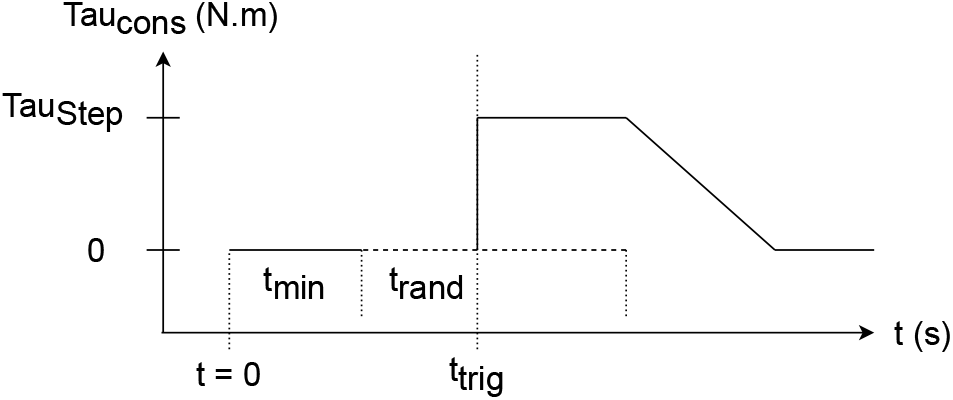
One jolt of torque

As a result of these tests, the user’s response time is measured with the following method. The maximum EMG while resting *U*_*rest,max,i*_ is recorded between *t* = 0 and *t*_*trig*_, where *i* denotes the EMG channel. *t*_*reac,i*_ is found with eq. (8) :

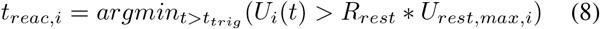

Where *R*_*rest*_ was set experimentally to 1.15. The user’s response time according to channel *i* is defined as : 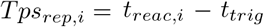. Finally, to reduce the variance, the result is averaged over the 3 channels placed over the biceps (as shown in eq. (9)) :

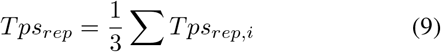

For the adaptation of the gain six jolts are applied in one position (3 upward, 3 downward) in order to keep the calibration step convenient.

### C. Experimental protocol

#### 1) Participants and equipement

The tests are conducted with 10 participants (height : 177, 2 ± 8, 4*cm*, weight : 77, 8 ± 20, 2*kg*, age : 24, 1 ± 1, 4). Each participant signed an informed consent before beginning the experiment. This protocol has been evaluated and validated through an ethical committee from Université Paris-Saclay (*n*° 212).

The exoskeleton used in this study is an under-actuated upper-body type (Fig. 8). Each side consists of two segments (upper-arm and forearm) and four joints. Two of the joints are passive (*θ*_1_ and *θ*_4_) and the other two are proportionally linked and powered by the same actuator (*θ*_3_ = 1.5∗*θ*_2_) [27](cf. Table II). The interface with the user consists in a rigid structure that is attached to the forearm, close to the wrist. It is attached with two pivot joints to end effector of the exoskeleton’s arm. In addition, the exoskeleton is backdriveable, which means that efforts applied to the end effector are transmitted to the actuator with minimum loss of energy [26].

**TABLE I:**
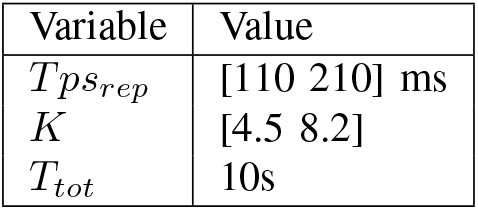
Range of Numerical values used in simulation

| Variable | Value |
| --- | --- |
| $Tps_{rep}$ | [110 210] ms |
| $K$ | [4.5 8.2] |
| $T_{tot}$ | 10s |

**TABLE II:** Table of the Denavit-Hartenberg parameters of the exoskeleton

| | $\alpha_i$ | $d_i$ | $r_i$ | $\theta_i$ |
| --- | --- | --- | --- | --- |
| Frame 1 | 0 | 0 | $r_{shoulder}$ | $\theta_1$ |
| Frame 2 | $+\frac{\pi}{2}$ | $d_{shoulder}$ | 0 | $\theta_2$ |
| Frame 3 | 0 | $L_{arm}$ | 0 | $\theta_3 = 1.5 * \theta_2$ |
| Frame 4 | $-\frac{\pi}{2}$ | $L_{elbow}$ | 0 | $\theta_4$ |
| Frame 5 | 0 | $L_{forearm}$ | 0 | 0 |

**TABLE III:**
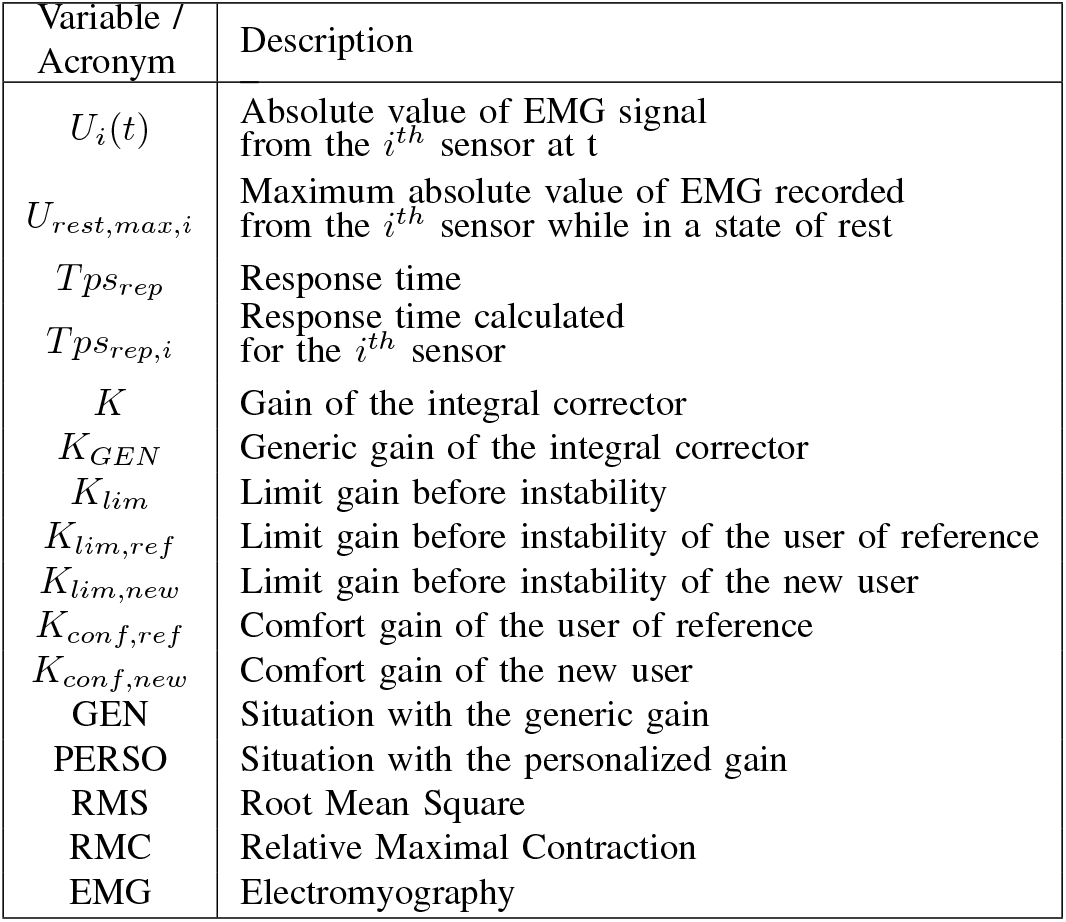
Nomenclature

**Fig. 8:**
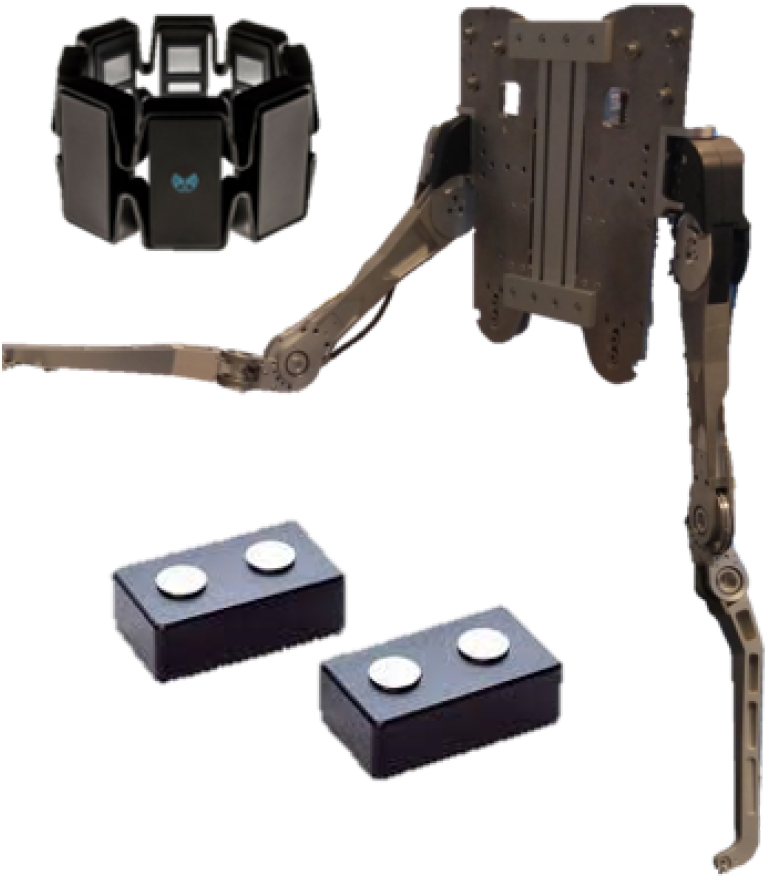
Upper-limb exoskeleton BHV2, Myo armband and Datalite EMG sensors

Two different types of EMG sensors are used in this study. The first one is the Myo-Armband (Thalmic Labs, ON, Canada), and is used to control the exoskeleton based on the work presented in [18] and [10]. The armband was positioned around the arm to capture biceps and triceps muscle activities, rather than on the foreman as it was originally designed. The Armband is composed of eight pairs of dry electrodes, with a sampling frequency of 200Hz. The raw output of the armband is a zero-mean signal coded over 8 bit, it has no unit and is comprised between -126 and +127. The second one are the sensors DataLite, Biometrics Ltd, Newport, UK. They are used to analyze muscles relevant during load carrying and have a sampling frequency of 1000Hz. The goal of this second type of measurement is to have an objective evaluation of the muscle activity during experimental conditions.

#### 2) Procedure

The participants are explained how to put on the EMG armband on the right arm. Then the EMG sensors are placed according the SENIAM recommendations [28] over the biceps, triceps, trapezius, anterior deltoid and erector spinae, on their left side.

Once equipped with the exoskeleton, the participants perform a lifting task in different situations. They start by doing 20 repetitions without assistance and with a load of 2kg, this is a reference situation. Then, they perform 50 repetitons with one of the two assistances in a randomized order and a 7-kilogram load. After this the participants are asked general questions about the assistance. They continue by doing the reference situation and the repetitions with the second assistance. Finally, they are asked general questions about the second assistance and questions of comparison between the two situations.

One repetition of the task is to lift a load up to a high mark, bring it down to a middle mark, high up again and finally put the load down (Fig.9). The reference repetitions enable to start the recordings of both situations in the same conditions. The targets are displayed on a screen as well as the current angular position (Fig.10). On this figure, the bottom and top black lines represent the exoskeleton’s joint limit. The blue and purple lines represent respectively the high and middle marks. The red line represents the angular position followed by the arm of the exoskeleton for one repetition of the task.

**Fig. 9:**
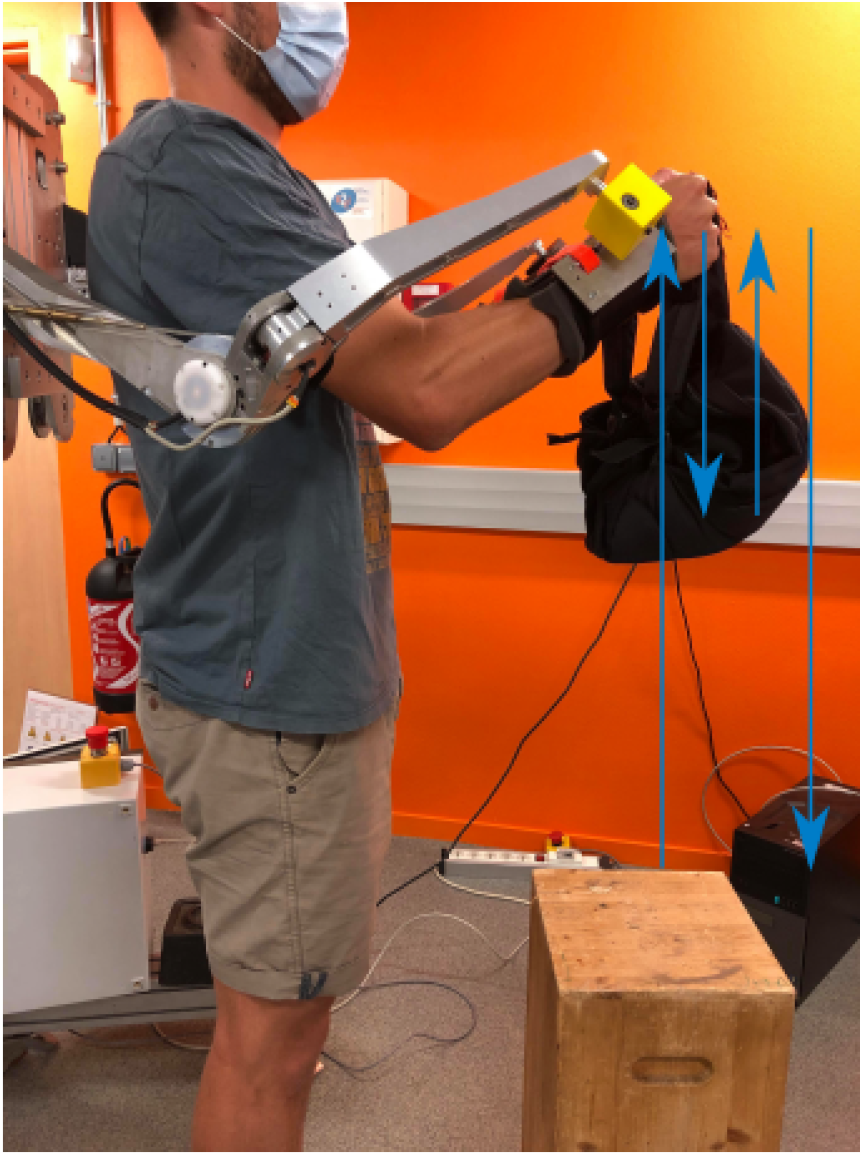
One repetition of the task

**Fig. 10:**
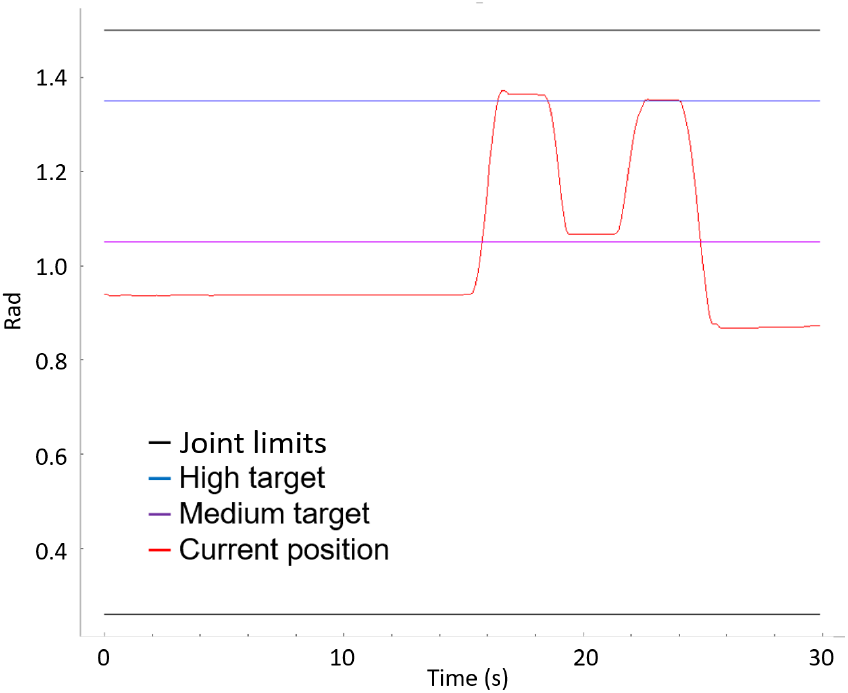
Display of the targets and current position

The difference between the two situations is the gain of the corrector, in one case it is generic (GEN), which means the same for each participant, in the other - it is personalized (PERSO), which means tuned following the method proposed in Methodology. The order was randomized, so 5 people started with GEN and 5 started with PERSO. When asking the question, the situations are referred to as “ Situation 1” and “Situation 2”, depending on the order and not the gain used, in order to avoid influencing the participants. Questionnaires are standard and the same for each situation. They have been designed according to the work of [29].

#### 3) Data Analysis

Three evaluation criteria are used during this study. The EMG signals and the precision are two objective criteria and the questionnaire is a subjective one.

Following the recommandations of [30] and [31], the EMG is first treated with a band pass Butterworth filter of order 7, with cutting frequencies [30Hz, 200Hz]. Then the signal is low-passed with a 7^*th*^ order Butterworth with a 2Hz cutting frequency [30]. In both cases the filter is double passed and the signals are normalized using the relative maximale contraction (RMC). The RMC is the maximum EMG value that occured for one muscle during the whole experiment (both tasks). The signal is then averaged over all repetitions and over series of 10 repetitions. The two situations and the evolution through the repetitions is evaluated with paired Student tests, with a threshold for significance of 0.05.

The precision is evaluated with the angular information of the exoskeleton’s shoulder. The participants are aksed to briefly stop at each target so the sequences where the velocity is near zero during the task are extracted. The error is calculted with the Root Mean Square (RMS) of the difference between the extracted positions and the targets positions. Similarly to the EMG signal, the averages for the whole 50 repetitions and series of 10 repetitions are calculated and compared with paired Student tests.

The questionnaire is divided in three parts : general questions about GEN, general questions about PERSO and questions about the comparison of the two. The answers take the form of a 11 point Likert scale (0 to 10) where 5 is the neutral value when it is releveant. The answers are averaged and the first two parts are compared through a Wilcoxon ranksign test. The same test is used to assess the shift from the neutral (5) of the comparison questions (last part).

## III. Results

### A. EMG

Fig. 11 shows the average muscle activity as a percentage of RMC during the task for the fifty repetitions. It is worth noting that there is a peak of biceps activity at the beginning of the task, engaging the assistance and a peak of triceps activity at the end, decreasing the assistance through the integrator. Then, the averages per series of 10 repetitions are calculated, in order to observe an evolution through time. It is represented on Fig. 12 for the anterior deltoid. A statistically significant reduction in mean activity is observed for both GEN and PERSO for the anterior deltoid between the first and the last series, *p <* 0.05. The activity goes from 28.6 ± 13.5% RMC to 17.2 ± 7.3% RMC for GEN and from 24.9 ± 8.5% RMC to 18.0 ± 6.8% RMC for PERSO. A similar evolution is observed for the trapezius but only significant for GEN, going from 22.1 ± 9.0% RMC to 14.7 ± 6.6% RMC. The other muscles (Biceps, Triceps and Erector Spinae) do not feature a significant change through the repetitions.

**Fig. 11:**
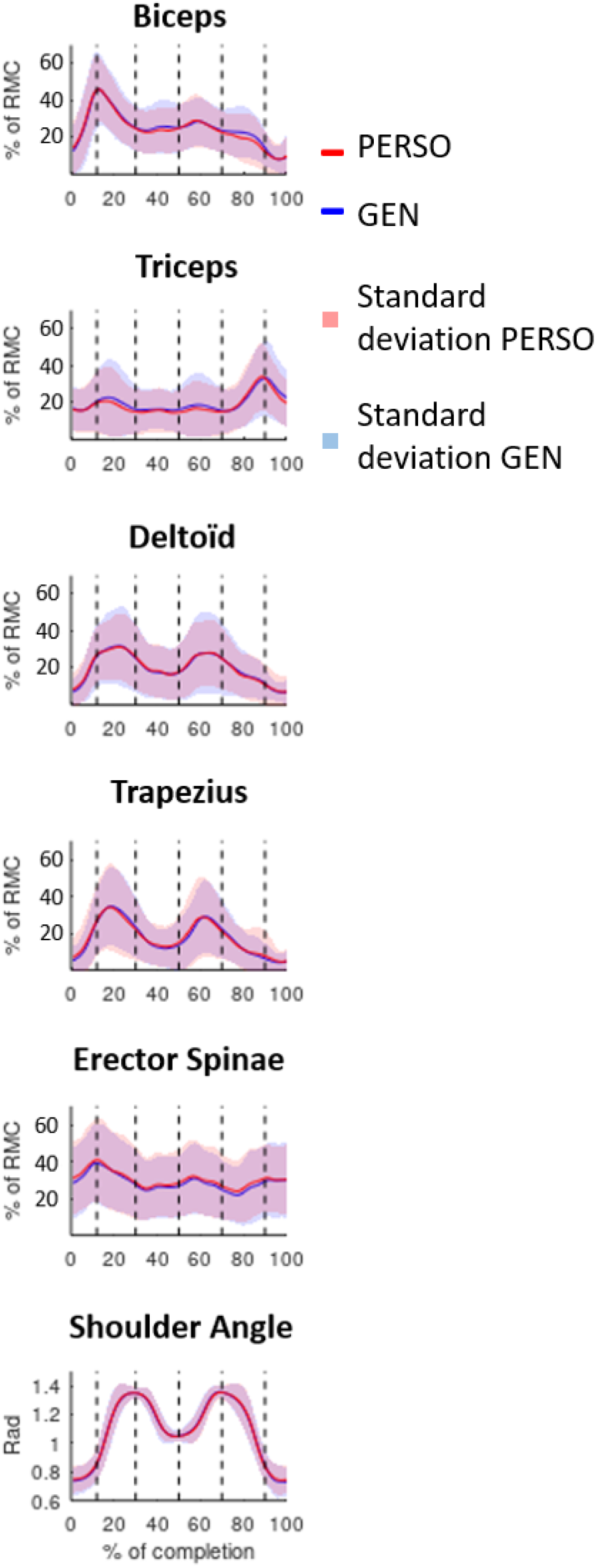
Average muscle activity and shoulder angle during one repetition (GEN in blue and PERSO in red)

**Fig. 12:**
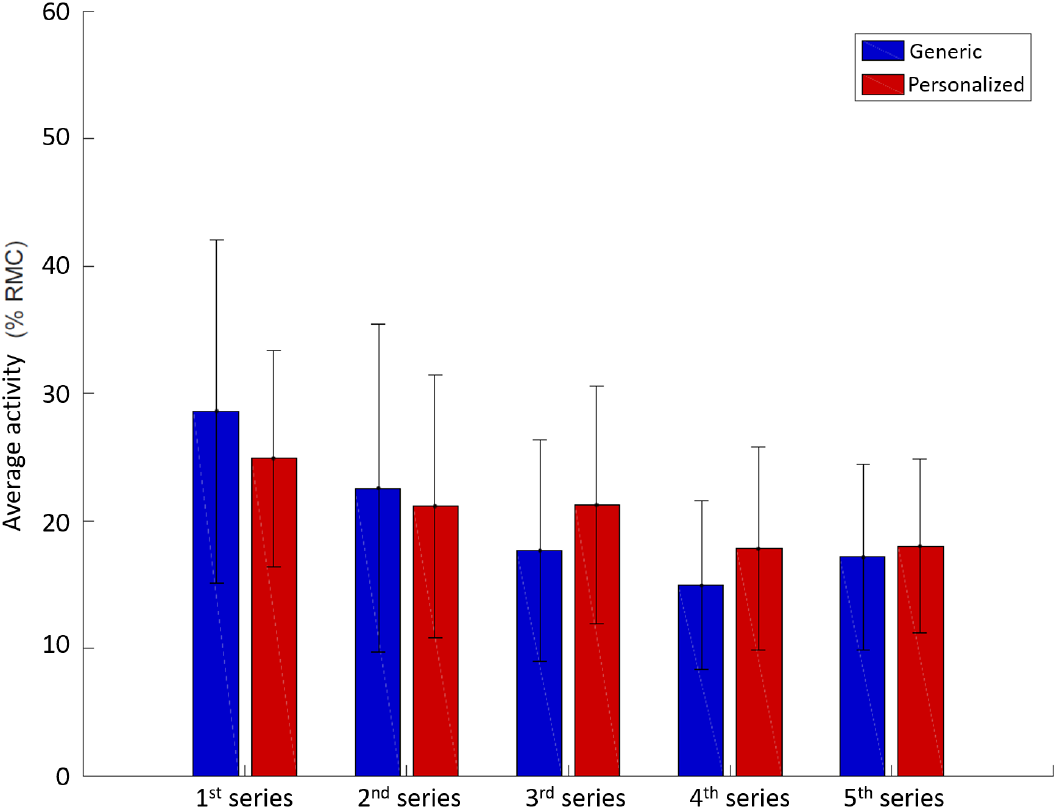
Average activity of the Deltoid per series of 10 repetitions (GEN in blue and PERSO in red)

The average activity of the anterior deltoid for the first series is displayed on Fig. 13. For 5 participants it is GEN and for the other 5 it is PERSO. The activity in the case of PERSO is significantly inferior (*p <* 0.05) to the one in the case of GEN (29.0 ± 8.0% RMC and 37.4 ± 9.5% RMC, respectively). However, this difference vanishes over the repetitions.

**Fig. 13:**
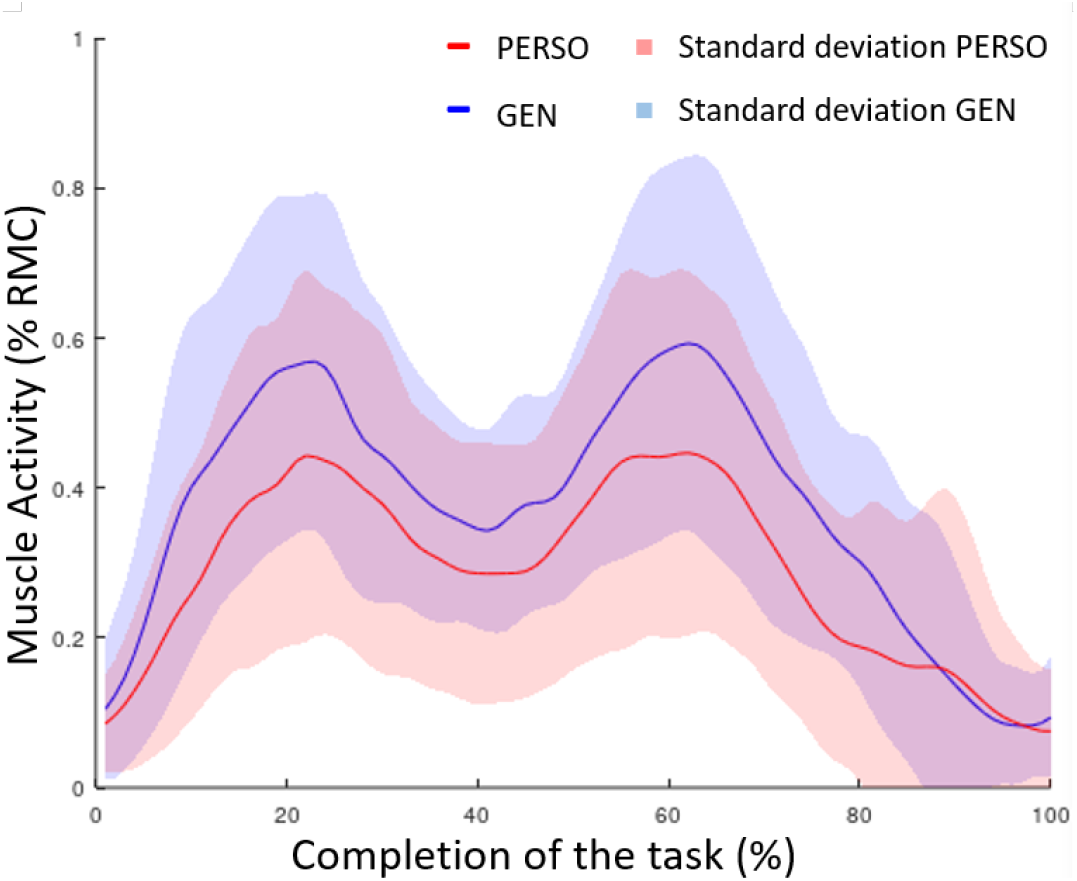
Average activity of the anterior deltoid for the first series (GEN in blue and PERSO in red)

### B. Precision

The precision achieved while aiming at the targets is averaged over the fifty repetitions in Fig. 14. Like with the EMG signal, both PERSO and GEN yield similar precision overall.

**Fig. 14:**
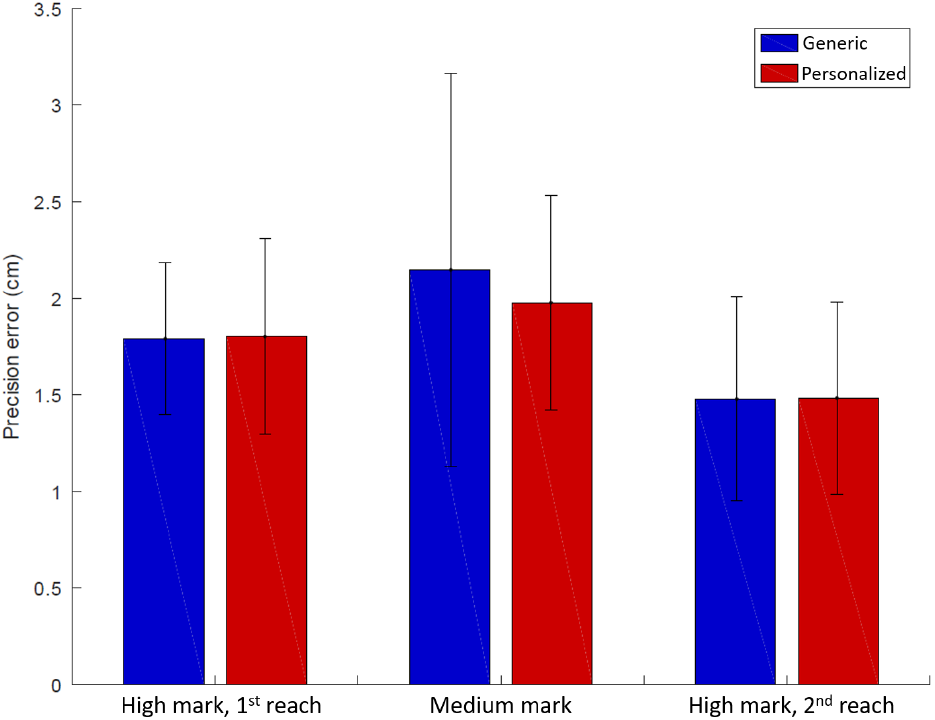
Average precision for each position

Similarly than with the EMG signal, the average precision per series of 10 repetitions for each key position (first high mark, middle mark and second high mark) is calculated. The results for the second reach of the high mark are represented on Fig. 15. There is an improvement of the precision for reaching the high target (both the first and second time), with an important reduction of the standard deviation. It goes from 2.02 ± 0.74 cm to 1.59 ± 0.33 cm for GEN and 2.29 ± 0.71 cm to 1.59 ± 0.29 cm for PERSO for the first reach of the high target. And it goes from 1.97 ± 1.14 cm to 1.27 ± 0.28 cm for GEN and 1.71 ± 0.87 cm to 1.26 ± 0.47 cm for PERSO for the second reach of the high target (cf. Fig. 15).

**Fig. 15:**
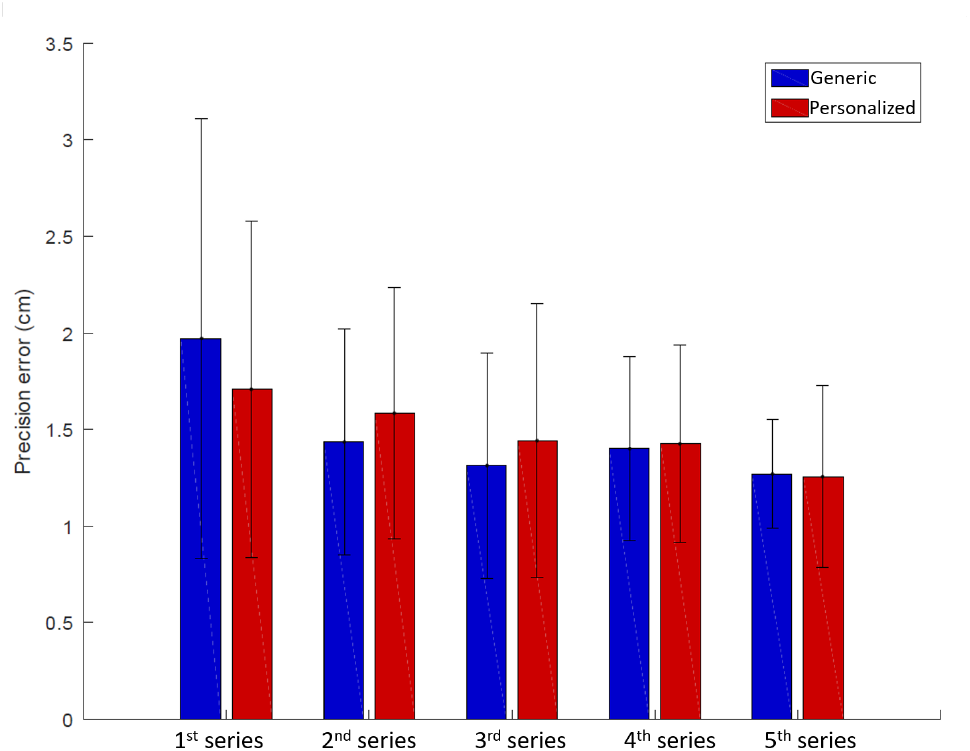
Precision for each series, for the high mark’s 2^*nd*^ reach

### C. Questionnaire

The answers to the general questions are moslty similar for the two situations as we can see on Fig. 16. The questions are divided in three categories : general, emotionnal affect and improvement over time (comparing the beginning and the end of a set). The “+” and “-” above the graph separate the positive key points from the negative ones. For example, a positive question is “I find it pleasant to work with the assistance” and a negative one is “Working with the assistance makes me anxious”. Overall, both assistances are well accepted, positive questions are above neutral on average and negative questions are around neutral or below. Only the first question about intuitiveness yields a statistically significant difference, according to the Wilcoxon test (*p <* 0.05). PERSO (7.5 ± 1.0) is perceived to be more intuitive than GEN (7.1 ± 0.7).

**Fig. 16:**
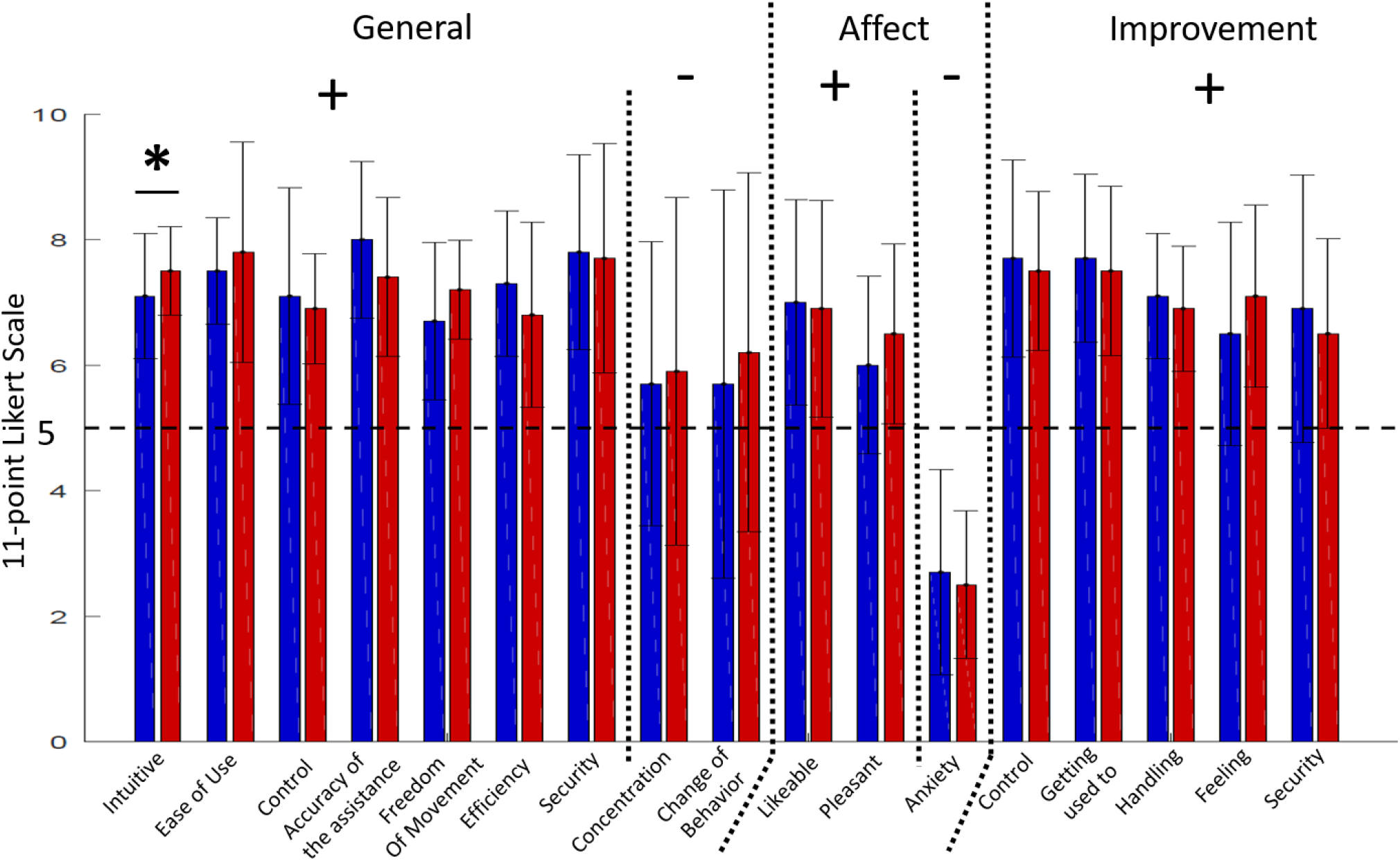
Average answers for the general questions (blue for GEN, red for PERSO)

The averaged answers to the comparison questions are displayed on Fig. 17. After a Wilcoxon test, a statistically significant shift of the results from the neutral value 5 is observed for 2 questions. The first one is about the intuitiveness and the second one is about the concentration required while performing the task. In both cases the shift advantages PERSO : it is judged more intuitive (5.8 ± 1.1) and requiring less concentration (4.5 ± 0.8).

**Fig. 17:**
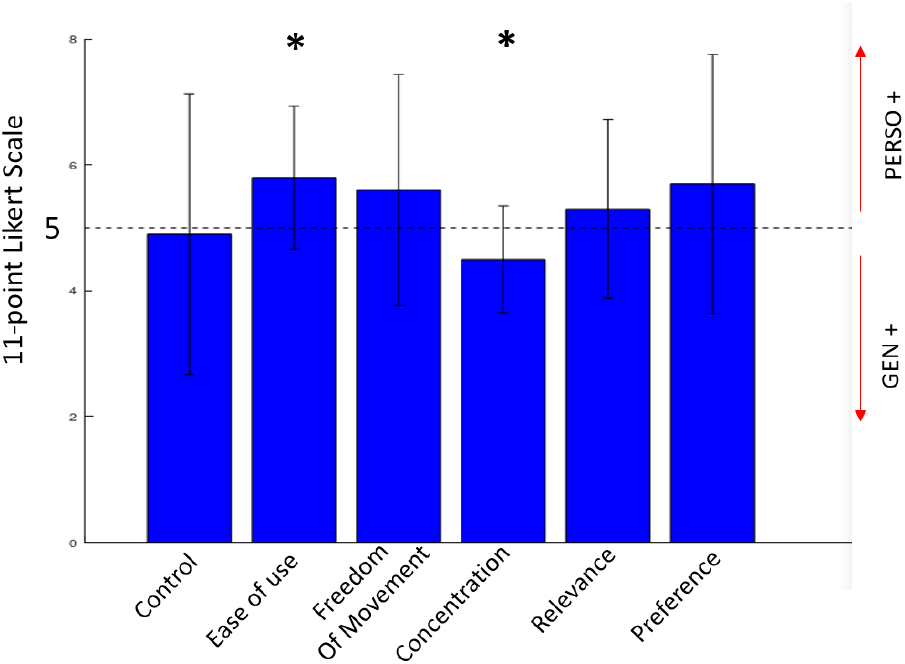
Average answers for the comparison questions

## IV. Discussion

The objectives of this study are to evaluate two points : (i) the impact of a proposed personalization of the control system of an exoskeleton and (ii) the adaptation of the user to the control system over several repetitions.

There are two main interpretations that can be made from the results presented in the previous section. The first one is that the proposed method of personalizing the gain improves the intuitiveness of the control system. Indeed, the post-processing of the EMG signals shows that the case of first contact presents a reduced activity of the anterior deltoid for PERSO (29.0 ± 8.0% RMC compared to 37.4 ± 9.5% RMC for GEN). These results align with the work of Yang et al. (2017), where the authors showed improved performances in teleoperation [32]. The approach designed by Yang and collaborators is a two step process : (i) user’s tremor attenuation based on a SVM and (ii) a real-time adaptation of the gain based on muscle activity. Authors show that their proposed method improve performances compared to conventional approaches. The improvement is caracterized by a reduced effort feedback and a reduced RMS error. In an other study Zhang and collaborators (2017) demonstrated the interest of human-in-the-loop optimization with an ankle exoskeleton [33]. These authors highlighted reduced metabolic energy cosumption with an optimized assistance compared to no assistance. This reduction is more important with an optimized assistance than with a generic one. Other related studies focused on the personalization of control systems. When considering gait rehabilitation, taking into account individual specificities is critical in order to provide relevant assistance [34], [35], [36]. The work of Chinimilli et al. (2019) showed a reduced muscular activity (vastus medialis) with a personalized walking assistance compared to baseline conditions. In addition to objective criteria our work also offered a subjective evaluation. The only general question that features a significant difference is about the intuiveness and the result is in favor of PERSO. Finally, when looking at the questions of comparison the same conclusion appears. Of the two questions that significantly shift from the neutral, one is about compared intuitiveness, favoring PERSO. The other one shows that the participants judged that PERSO requires less concentration. This further reinforces the point that PERSO is more intuitive since an intuitive system should not require more concentration to be used.

The second interpretation is that it is not necessary to personalize the system to ensure a safe and efficient interaction. The results obtained from the EMG signal show that both situations feature a reduction of activity of the anterior deltoid along the repetitions. In the end, the activation levels are similar for both situations. The same type of reduction occurs for the trapezius for GEN as well. However, this decrease does not occur for the trapezius muscle in the PERSO condition. This may be due to the fact that initial muscle activity of trapezius in this condition is lower than in the GEN condition. In addition, precision in reaching the targets improves in a similar way for both situations. The results of the general questions show that both situations performed well concerning the acceptance of the user. These positive elements, i.e. precision and acceptance, support the idea of a generic gain that might be deduced from simulation. Indeed, modelling the human behavior permits to define a range of values for the gain with boundary conditions (see Fig. 6). The shorter the response time, the greater can be the gain. On the contrary, if we consider a maximum response time that we expect to see in a given population, we can pick a value for the gain that would ensure the stability for every member of this group. The gain would not be the most suitable for everyone but results mentionned earlier suggest that users would adapt anyway. This interpretation also align with the work of Zhang et al. where they show that a generic assistance is also beneficial (albeit not as efficient as a personalized one) [33].

## V. Conclusion

In this study a method to adapt the gain of an integral-based control system of an exoskeleton to improve user’s intuitiveness was presented. The adaptation was based on an objective criteria, and simulations were used to determine the relation between gain and response time. An experiment was conducted with the help of 10 participants in order to compare the personalized gain to a generic one.

The main contribution of this study was the evaluation of the impact of the proposed personalization method rather than only the personalization itself. Indeed, a system of assistance that was shown to work in a generic way was chosen [10] and an automated approach was proposed to personalize the main parameter of control, which was the gain of the integral corrector.

Results showed that personalizing the gain improved the intuiteveness of the system, but also that with some training both gains yielded similar results. A complementary study might be conducted in order to let participants select the gain they prefer, while still measuring their response time. This alternative approach could potentially confirm the proposed method of adaptation, or offer an empirical relation between gain and response time.

## References

[1] Nicolas Vignais, Markus Miezal, Gabriele Bleser, Katharina Mura, Dominic Gorecky, and Frédéric Marin. Innovative system for real-time ergonomic feedback in industrial manufacturing. Applied ergonomics, 44(4):566–574, 2013.

[2] INRS. Troubles musculo-squelettiques - statistiques. 2015.

[3] L’Assurance Maladie. Risques professionnels : Rapport annuel, 2017.

[4] Philippe Garrec. Design of an anthropomorphic upper limb exoskeleton actuated by ball-screws and cables. Bulletin of the Academy of Sciences of the Ussr-Physical Series, 72(2):23, 2010.

[5] JianTao Yang and Cheng Peng. Adaptive motion intent understanding– based control of human–exoskeleton system. Proceedings of the Institution of Mechanical Engineers, Part I: Journal of Systems and Control Engineering, page 0959651820945814, 2020.

[6] Dongil Choi and Jun-ho Oh. Development of the cartesian arm exoskeleton system (caes) using a 3-axis force/torque sensor. International Journal of Control, Automation and Systems, 11(5):976–983, 2013.

[7] Christian Fleischer and Günter Hommel. A human–exoskeleton interface utilizing electromyography. IEEE Transactions on Robotics, 24(4):872–882, 2008.

[8] Stefano Toxiri, Axel S Koopman, Maria Lazzaroni, Jesús Ortiz, Valerie Power, Michiel P de Looze, Leonard O’Sullivan, and Darwin G Caldwell. Rationale, implementation and evaluation of assistive strategies for an active back-support exoskeleton. Frontiers in Robotics and AI, 5:53, 2018.

[9] Luka Peternel, Tomoyuki Noda, Tadej Petrič, Aleš Ude, Jun Morimoto, and Jan Babič. Adaptive control of exoskeleton robots for periodic assistive behaviours based on emg feedback minimisation. PloS one, 11(2):e0148942, 2016.

[10] Benjamin Treussart, Franck Geffard, Nicolas Vignais, and Frédéric Marin. Controlling an upper-limb exoskeleton by emg signal while carrying unknown load. In 2020 IEEE International Conference on Robotics and Automation (ICRA), pages 9107–9113. IEEE, 2020.

[11] Arash Ajoudani, Andrea Maria Zanchettin, Serena Ivaldi, Alin Albu-Schäffer, Kazuhiro Kosuge, and Oussama Khatib. Progress and prospects of the human–robot collaboration. Autonomous Robots, pages 1–19, 2018.

[12] C DaSalla, J Kim, and Y Koike. Robot control using electromyography (emg) signals of the wrist. Applied Bionics and Biomechanics, 2(2):97–102, 2005.

[13] Pornchai Phukpattaranont, Sirinee Thongpanja, Khairul Anam, Adel Al-Jumaily, and Chusak Limsakul. Evaluation of feature extraction techniques and classifiers for finger movement recognition using surface electromyography signal. Medical & biological engineering & computing, pages 1–13, 2018.

[14] Marta Gandolla, Simona Ferrante, Giancarlo Ferrigno, Davide Baldassini, Franco Molteni, Eleonora Guanziroli, Michele Cotti Cottini, Carlo Seneci, and Alessandra Pedrocchi. Artificial neural network emg classifier for functional hand grasp movements prediction. Journal of International Medical Research, 45(6):1831–1847, 2017.

[15] Felix E Zajac. Muscle and tendon properties models scaling and application to biomechanics and motor. Critical reviews in biomedical engineering, 17(4):359–411, 1989.

[16] Panagiotis K Artemiadis and Kostas J Kyriakopoulos. An emg-based robot control scheme robust to time-varying emg signal features. IEEE Transactions on Information Technology in Biomedicine, 14(3):582–588, 2010.

[17] Sebastian Amsuess, Ivan Vujaklija, Peter Goebel, Aidan D Roche, Bernhard Graimann, Oskar C Aszmann, and Dario Farina. Contextdependent upper limb prosthesis control for natural and robust use. IEEE Transactions on Neural Systems and Rehabilitation Engineering, 24(7):744–753, 2015.

[18] Benjamin Treussart, Franck Geffard, Nicolas Vignais, and Frédéric Marin. Controlling an exoskeleton with emg signal to assist load carrying: a personalized calibration. In 2019 International Conference on Mechatronics, Robotics and Systems Engineering (MoRSE), pages 246–252. IEEE, 2019.

[19] JR Potvin and SHM Brown. Less is more: high pass filtering, to remove up to 99% of the surface emg signal power, improves emg-based biceps brachii muscle force estimates. Journal of Electromyography and Kinesiology, 14(3):389–399, 2004.

[20] Khalil Ullah and Jung-Hoon Kim. A mathematical model for mapping emg signal to joint torque for the human elbow joint using nonlinear regression. In ICARA 2009, pages 103–108. IEEE, 2009.

[21] Karen Simonyan and Andrew Zisserman. Very deep convolutional networks for large-scale image recognition. arXiv preprint 1409.1556, 2014.

[22] Stephen H Scott. Optimal feedback control and the neural basis of volitional motor control. Nature Reviews Neuroscience, 5(7):532–545, 2004.

[23] Tamar Flash and Neville Hogan. The coordination of arm movements: an experimentally confirmed mathematical model. Journal of neuroscience, 5(7):1688–1703, 1985.

[24] Simon Bastide, Nicolas Vignais, Franck Geffard, and Bastien Berret. Interacting with a “transparent” upper-limb exoskeleton: A human motor control approach. In 2018 IEEE/RSJ International Conference on Intelligent Robots and Systems (IROS), pages 4661–4666. IEEE, 2018.

[25] DFB Haeufle, M Günther, A Bayer, and S Schmitt. Hill-type muscle model with serial damping and eccentric force–velocity relation. Journal of biomechanics, 47(6):1531–1536, 2014.

[26] Tatsuzo Ishida and Atsuo Takanishi. A robot actuator development with high backdrivability. In Robotics, Automation and Mechatronics, 2006 IEEE Conference on, pages 1–6. IEEE, 2006.

[27] Philippe Garrec, Yann Perrot, Dominique Ponsort, and Aurelie Riglet. Patent: Exoskeleton arm having an actuator. 2012. US9375325B2.

[28] Hermie J Hermens, Bart Freriks, Roberto Merletti, Dick Stegeman, Joleen Blok, Günter Rau, Cathy Disselhorst-Klug, and Göran Hägg. European recommendations for surface electromyography. Roessingh research and development, 8(2):13–54, 1999.

[29] P. Maurice, J. Čamernik, D. Gorjan, B. Schirrmeister, J. Bornmann, L. Tagliapietra, C. Latella, D. Pucci, L. Fritzsche, S. Ivaldi, and J. Babič. Objective and subjective effects of a passive exoskeleton on overhead work. IEEE Transactions on Neural Systems and Rehabilitation Engineering, 28(1):152–164, 2020.

[30] Thomas S Buchanan, David G Lloyd, Kurt Manal, and Thor F Besier. Neuromusculoskeletal modeling: estimation of muscle forces and joint moments and movements from measurements of neural command. Journal of applied biomechanics, 20(4):367–395, 2004.

[31] Alison C McDonald, Kia Sanei, and Peter J Keir. The effect of high pass filtering and non-linear normalization on the emg–force relationship during sub-maximal finger exertions. Journal of Electromyography and Kinesiology, 23(3):564–571, 2013.

[32] Chenguang Yang, Jing Luo, Yongping Pan, Zhi Liu, and Chun-Yi Su. Personalized variable gain control with tremor attenuation for robot teleoperation. IEEE Transactions on Systems, Man, and Cybernetics: Systems, 48(10):1759–1770, 2017.

[33] Juanjuan Zhang, Pieter Fiers, Kirby A Witte, Rachel W Jackson, Katherine L Poggensee, Christopher G Atkeson, and Steven H Collins. Human-in-the-loop optimization of exoskeleton assistance during walking. Science, 356(6344):1280–1284, 2017.

[34] L. Rose, M. C. F. Bazzocchi, and G. Nejat. End-to-end deep reinforcement learning for exoskeleton control. In 2020 IEEE International Conference on Systems, Man, and Cybernetics (SMC), pages 4294–4301, 2020.

[35] Prudhvi Tej Chinimilli, Zhi Qiao, Seyed Mostafa Rezayat Sorkhabadi, Vaibhav Jhawar, Iat Hou Fong, and Wenlong Zhang. Automatic virtual impedance adaptation of a knee exoskeleton for personalized walking assistance. Robotics and Autonomous Systems, 114:66–76, 2019.

[36] Manuel Cardona, Cecilia E Garcia Cena, Fernando Serrano, and Roque Saltaren. Alice: conceptual development of a lower limb exoskeleton robot driven by an on-board musculoskeletal simulator. Sensors, 20(3):789, 2020.

